# Quality control implementation for universal characterization of DNA and RNA viruses in clinical respiratory samples using single metagenomic next-generation sequencing workflow

**DOI:** 10.1101/367367

**Authors:** A. Bal, M. Pichon, C. Picard, JS. Casalegno, M. Valette, I. Schuffenecker, L. Billard, S. Vallet, G. Vilchez, V. Cheynet, G. Oriol, S. Assant, Y. Gillet, B. Lina, K. Brengel-Pesce, F. Morfin, L. Josset

**Affiliations:** Laboratoire de Virologie, Institut des Agents Infectieux, Groupement Hospitalier Nord, Hospices Civils de Lyon, Lyon, France; Univ Lyon, Université Lyon 1, Faculté de Médecine Lyon Est, CIRI, Inserm U1111 CNRS UMR5308, Virpath, Lyon, France; Hospices Civils de Lyon, Centre National de Reference des virus respiratoires France Sud, Lyon, France; Laboratoire Commun de Recherche HCL-bioMerieux, Centre Hospitalier Lyon Sud, Pierre-Bénite, France; Unité de Biologie des Infections Virales Emergentes, Institut Pasteur, Lyon, France; CIRI Inserm U1111, CNRS 5308, ENS, UCBL, Faculté de Médecine Lyon Est, Université de Lyon, Lyon, France; INSERM UMR1078 “Génétique, Génomique Fonctionnelle et Biotechnologies”, Axe Microbiota, Univ Brest, Brest, France; Département de Bactériologie-Virologie, Hygiène et Parasitologie-Mycologie, Pôle de Biologie-Pathologie, Centre Hospitalier Régional et Universitaire de Brest, Hôpital de la Cavale Blanche, Brest, France; Hospices Civils de Lyon, Urgences pédiatriques, Hôpital Femme Mère Enfant, Bron, France

**Keywords:** clinical virology, quality control, next-generation sequencing, viral metagenomics, respiratory viruses

## Abstract

**Background:** In recent years, metagenomic Next-Generation Sequencing (mNGS) has increasingly been used for an accurate assumption-free virological diagnosis. However, the systematic workflow evaluation on clinical respiratory samples and implementation of quality controls (QCs) is still lacking.

**Methods:** A total of 3 QCs were implemented and processed through the whole mNGS workflow: a notemplate-control to evaluate contamination issues during the process; an internal and an external QC to check the integrity of the reagents, equipment, the presence of inhibitors, and to allow the validation of results for each sample. The workflow was then evaluated on 37 clinical respiratory samples from patients with acute respiratory infections previously tested for a broad panel of viruses using semi-quantitative real-time PCR assays (28 positive samples including 6 multiple viral infections; 9 negative samples). Selected specimens included nasopharyngeal swabs (n = 20), aspirates (n = 10), or sputums (n = 7).

**Results:** The optimal spiking level of the internal QC was first determined in order to be sufficiently detected without overconsumption of sequencing reads. According to QC validation criteria, mNGS results were validated for 34/37 selected samples. For valid samples, viral genotypes were accurately determined for 36/36 viruses detected with PCR (viral genome coverage ranged from 0.6% to 100%, median = 67.7%). This mNGS workflow allowed the detection of DNA and RNA viruses up to a semi-quantitative PCR Ct value of 36. The six multiple viral infections involving 2 to 4 viruses were also fully characterized. A strong correlation between results of mNGS and real-time PCR was obtained for each type of viral genome (R^2^ ranged from 0.72 for linear single-stranded (ss) RNA viruses to 0.98 for linear ssDNA viruses).

**Conclusions:** Although the potential of mNGS technology is very promising, further evaluation studies are urgently needed for its routine clinical use within a reasonable timeframe. The approach described herein is crucial to bring standardization and to ensure the quality of the generated sequences in clinical setting. We provide an easy-to-use single protocol successfully evaluated for the characterization of a broad and representative panel of DNA and RNA respiratory viruses in various types of clinical samples.

## Background

Since the development of Next Generation-Sequencing (NGS) technologies in 2005, the use of metagenomic approaches has grown considerably. It is now considered as an efficient unbiased tool in clinical virology[1,2], in particular for the characterization of viral acute respiratory infections (ARIs). Several advantages of metagenomic NGS (mNGS) compared to conventional real-time Polymerase Chain Reaction (PCR) assays have been highlighted. Firstly, the full viral genetic information is immediately available allowing the investigation of respiratory outbreaks, viral epidemiological surveillance, or identification of specific mutations leading to antiviral resistance or higher virulence [3–5]. Secondly, a significant improvement in viral ARIs diagnosis has been reported [4,6–9]; as the process is sequence independent, mNGS is able to identify highly divergent viral genomes, rare respiratory pathogens, and to discover respiratory viruses missed by targeted PCR [1,4,7].

However, the diversity in viral nucleic acid types has impaired the development of a unique viral metagenomic workflow allowing the comprehensive characterization of viruses present in a clinical sample. Most of the published viral metagenomic protocols have been optimized for the detection either of DNA viruses or RNA viruses [4,5,10–13]. In addition, despite the growing number of studies using a metagenomic process in clinical virology, evaluation of workflows has not systematically included both clinical samples and quality control (QC) implementation. A metagenomic protocol involves a large number of steps and all of these have to be controlled to ensure the quality of the generated sequences [6,14–16]. Furthermore, specimen to specimen, environmental, and reagent contaminations are also a major concern in metagenomic setting and must be accurately evaluated [6,17–19].

The objective of this study was to implement QCs in a single metagenomic protocol and to evaluate it for the detection of a broad panel of DNA and RNA viruses in clinical respiratory samples.

## Methods

### Clinical samples

A total of 37 respiratory samples collected from patients hospitalized in the university hospital of Lyon (Hospices Civils de Lyon, HCL) were retrospectively selected to evaluate our metagenomic approach. Selected specimens included various types of clinical samples; nasopharyngeal swabs (n=20), aspirates (n=10), or sputums (n=7). These samples were initially sent to our laboratory for routine viral diagnosis of ARI using semi-quantitative real-time PCR assays targeting a comprehensive panel of DNA and RNA viruses (r-gene, bioMérieux, Marcy l’étoile, France). This panel included: influenza virus type A and B, adenovirus, cytomegalovirus, Epstein-Barr virus, human herpes virus 6, human bocavirus (HBoV), human rhinovirus, respiratory syncytial virus, human parainfluenza virus, human coronavirus (HCoV), human metapneumovirus, and measles virus. Twenty-two samples were positive for only one targeted virus, 6 were characterized by a multiple viral infection and 9 were negative for all the targeted viruses. These 9 samples were also found to be negative using the FilmArray Respiratory Panel (FA RP, bioMérieux). After PCR testing, the rest of samples were stored at –20°C until mNGS analysis.

### Metagenomic workflow

For sample viral enrichment, a 3-step method was applied to 200μl of thawed and vortexed sample [20]: low-speed centrifugation (6000g, 10 min, 4 °C), followed by filtration of the supernatant using 0.80 µm filter (Sartorius, Göttingen, Germany) to remove eukaryotic and bacterial cells, without loss of large viruses [21] and then Turbo DNase treatment (0.1U/μL, 37 °C, 90 min; Life Technologies, Carlsbad, CA, USA). Total nucleic acid was extracted using the NucliSENS EasyMAG platform (bioMérieux, Marcy l’Etoile, France) followed by an ethanol precipitation (2 hours at –80°C). As previously described, modified whole transcriptome amplification was performed to amplify both DNA and RNA viral nucleic acids (WTA2, Sigma-Aldrich, Darmstadt, Germany) [21]. Amplified DNA and cDNA were then purified using a QiaQuick column (Qiagen, Hilden, Germany) and quantified using the Qubit fluorometer HS dsDNA Kit (Life Technologies, Carlsbad, CA, USA). Nextera XT DNA Library preparation and Nextera XT Index Kit were used to prepare paired-end libraries, according to the manufacturer’s recommendations (Illumina, San Diego, CA, USA). After normalization, a pool of libraries (V/V) was made and quantified using universal KAPA library quantification kit (Kapa Biosystems, Wilmington, MA, USA); 1% PhiX genome was added to the quantified library before sequencing with Illumina NextSeq 500 ™ platform (Fig. 1). In addition, it should be noticed that our wet-lab process was designed to prevent contaminations as much as possible: reagents were stored and prepared in a DNA-free room; patient samples were opened in a laminar flow hood in a pre-PCR room; after the amplification step, tubes were handled and stored in a post-PCR room.

**Fig. 1.**
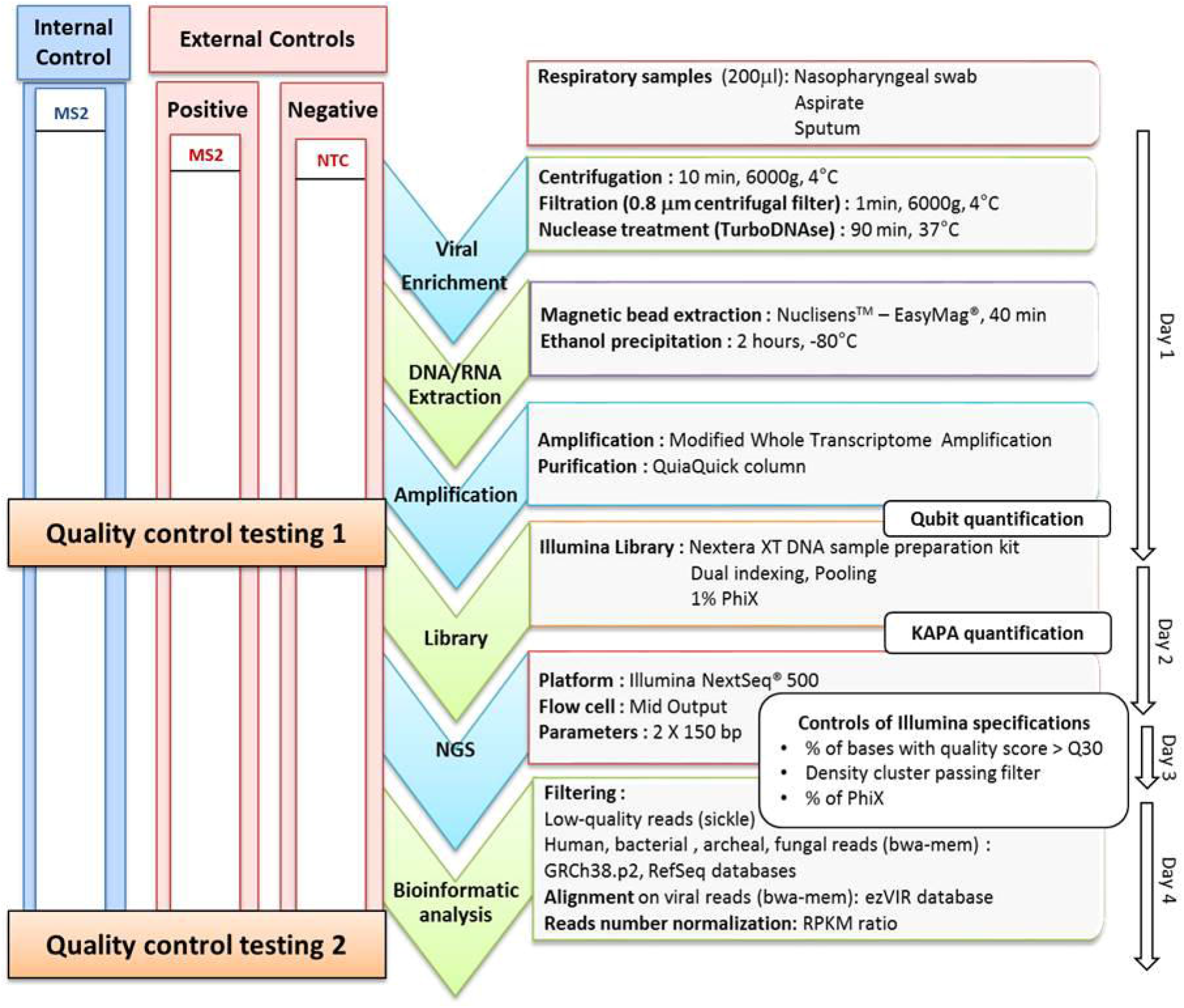
Schematic representation of the metagenomic workflow and quality control steps. The whole process is summarized in the middle. On the left side, internal control (MS2 bacteriophage) is represented in blue, and external controls are represented in red, including positive control (MS2 bacteriophage spiked in viral transport medium) and No-Template Control (NTC: viral transport medium). Quality control testing 1 corresponds to MS2 bacteriophage molecular detection with commercial PCR assay. Quality control testing 2 corresponds to control by sequencing metrics (number of MS2 reads normalized with RPKM ratio and MS2 genome coverage). On the right, each technique used by phases is indicated black. In addition, on the far right the duration of each step is indicated.

### Bioinformatic analysis

A stepwise bioinformatic filtering pipeline was used to quality filter reads using cutadapt and sickle; and to remove human, archaeal, bacterial, and fungal sequences by aligning reads with bwa mem. The databases used were GRCh38.p2, RefSeq archaea, RefSeq bacteria, and RefSeq fungi. Remaining reads were aligned on ezVIR viral database v0.1 [22] and bacteriophage genomes from the RefSeq database (downloaded on 17 February 2017) using bwa mem. Normalization for comparing viral genome coverage values was performed using reads per kilobase of virus reference sequence per million mapped reads (RPKM) ratio [4,23]. RPKM ratio corrects differences in both sample sequencing depth and viral gene length. Viral reads (expressed in RPKM) from the No-Template Control (NTC) were subtracted from viral reads (in RPKM) of each sample within the batch prior to further analysis. A sample was considered to be positive for a particular virus when the RPKM of this virus was positive. No threshold regarding genome coverage pattern was applied nor requirement to cover a particular region of the genome. This latter requirement could be important to correctly identify RNA virus subtypes with high recombination frequencies within a species, but has to be implemented specifically for each viral family.

### Quality control implementation

All respiratory specimens were spiked with internal quality control (IQC) before sample preparation. MS2 bacteriophage from a commercial kit (MS2, IC1 RNA internal control; r-gene, bioMérieux) was selected as the IQC. As positive external quality control (EQC), we used viral transport medium spiked with MS2 at the same concentration used for the IQC. A No-Template Control (NTC) was implemented to evaluate contamination during the process. NTC was constituted of viral transport medium (Sigma-virocult, MWE, Corsham, UK) that was processed through all mNGS steps. Two QC testing (QCT) were performed: QCT1 which was the semi-quantitative detection of MS2 using a commercial real-time PCR assay (IC1 RNA internal control, r-gene, bioMérieux,) after amplification step (Fig. 1). QCT1 validation criteria were: MS2 semi-quantitative PCR Cycle threshold (Ct) below 37 Ct for IQC and EQC, and no MS2 detection for NTC. QCT2 evaluated the sequencing performance by quantifying the number of reads aligned on the MS2 genome (in RPKM) and MS2 genome coverage (MS2 genome accession number: NC_001417.2; Fig. 1). QCT2 validation criteria were MS2 genome coverage >95% for positive EQC, and an MS2 RPKM > 0 for IQC.

### Statistical analysis

Statistical analyses were performed using GraphPrism version 5.02 applying the appropriate statistical test (associations between mNGS and viral real-time PCR assay were determined by applying the Pearson’s correlation coefficient and differences between median and distributions were evaluated by the Mann–Whitney U test). A p-value less than 0.05 was considered to be statistically significant.

## Results

### Determination of optimal internal quality control spiking

MS2 bacteriophage (MS2), a single-stranded RNA virus (ssRNA), was used as the IQC to validate the whole metagenomic process for each sample. In order to optimize IQC spiking level, the sensitivity of the metagenomic analysis workflow for MS2 detection was first evaluated with a ten-fold serial dilutions of MS2 (from 10^−2^ to 10^−5^) in a nasopharyngeal swab tested negative using FA RP (bioMérieux). MS2 was detected in internal QCT1 (IQCT1) for all levels of MS2 spiking (Ct ranged from 17.5 at the 10^−2^ dilution to 26.4 Ct at the 10^−5^ dilution). Full to partial MS2 genome coverage was obtained for all MS2 spiking levels in internal QCT2 (IQCT2; coverage ranged from 98% at the 10^−2^ dilution to 69% at the 10^−5^ dilution). For the highest spiking level, 66.0% of the total number of viral reads was mapped to MS2; for the lowest spiking level, 0.9% were so (Fig. 2). To limit the number of NGS reads consumed for IQC detection, the optimal spiking condition was determined to be the 10^−5^ dilution and was used for the rest of the study.

**Fig. 2.**
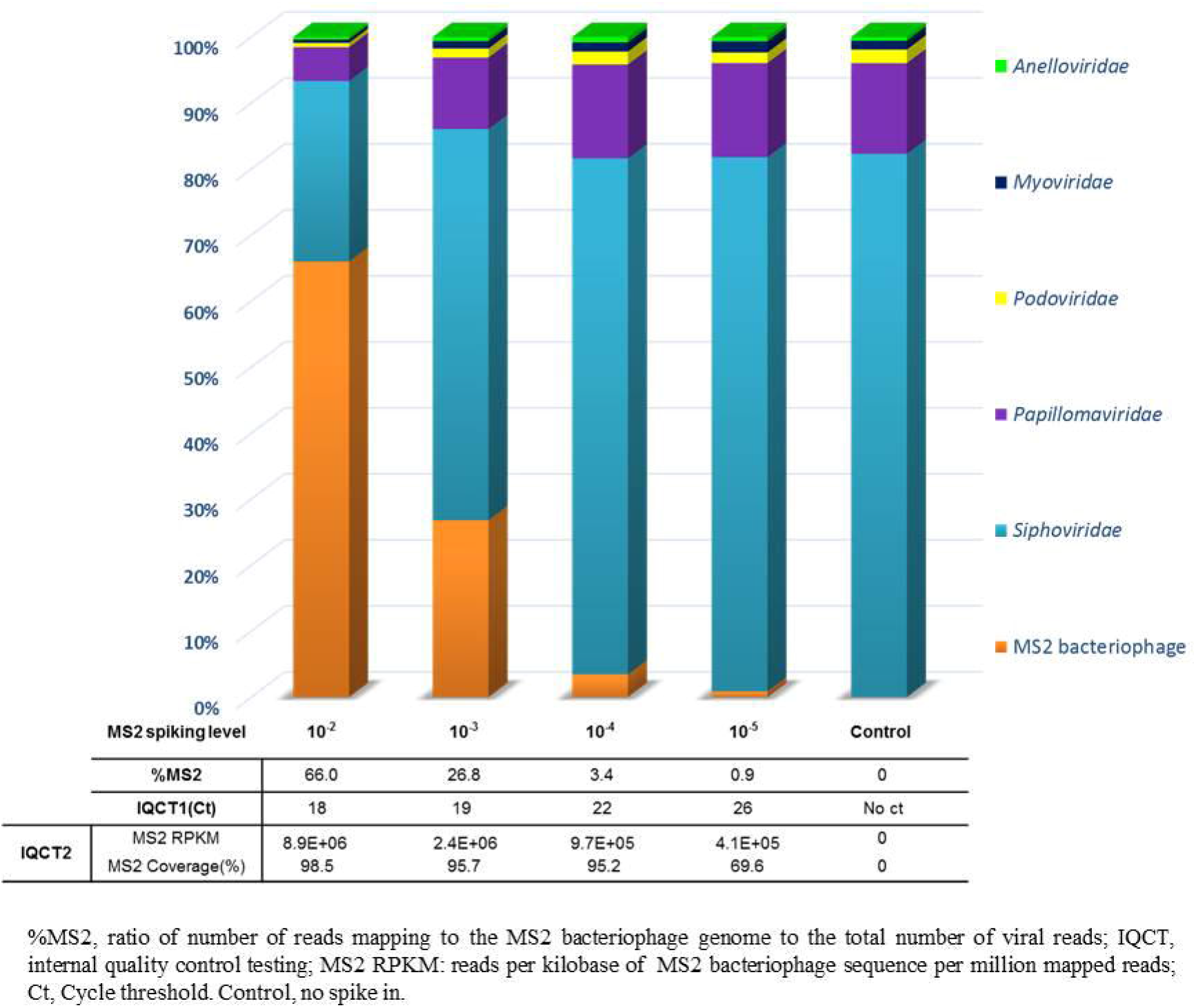
Determination of optimal spiking level for internal quality control. The sensitivity of the metagenomic analysis workflow for MS2 bacteriophage (Internal Quality Control, IQC) detection was evaluated with a MS2 ten-fold serial dilutions in a nasopharyngeal swab tested negative with multiplex viral PCR. Relative abundance of MS2 bacteriophage and viral families are represented depending on the MS2 spiked-in concentration. IQCT1 corresponds to MS2 molecular detection with commercial real-time PCR assay after amplification step. IQCT2 corresponds to control by sequencing metrics (number of MS2 reads normalized with RPKM ratio and MS2 genome coverage).

### Validation of mNGS results

A total of 37 clinical respiratory samples from patients with ARIs caused by a broad panel of DNA and RNA viruses or of unknown etiology were analyzed in a single mNGS workflow. Libraries were sequenced to a mean of 5,139,248 million reads passing quality filters (range: 270,975 to 13,586,456 reads). Human sequences represented the main part of NGS reads for both positive samples (mean = 61.3%) and negative samples (mean = 67.1%), but not of NTC which was mainly composed of bacterial reads (67.8%). The proportion of viral reads ranged from 0.006% to 85.2% (mean = 9.6% for positive samples and 0.6 % for negative samples, Additional file 1). Viral metagenomic results were then validated according to the criteria described in the Methods section. QCT1 (MS2 molecular detection performed before library preparation) was negative for NTC. After sequencing, viral contamination represented 0.13% (4,245/3,215,616) of the total reads generated from NTC including 2 MS2 reads (MS2 RPKM = 173). For targeted viruses, 21 reads (RPKM = 480) and 185 reads (RPKM = 1.1E+04) mapping to influenza A(H3N2) and HBoV were detected, respectively. The positive EQC was successfully detected at QCT1 (MS2 PCR positive at 25 Ct) and after the sequencing step (QCT2; MS2 genome coverage = 99.7%, MS2 RPKM = 5.5E+05). Regarding IQC results, 37/37 samples passed QCT1 (MS2 PCR Ct values <37) and were therefore further processed. A total of 33/37 samples passed QCT2 (MS2 RPKM > 0; Fig. 3). For these 33 samples, MS2 genome coverage ranged from 15% to 100% (Additional file 2).

**Fig. 3.**
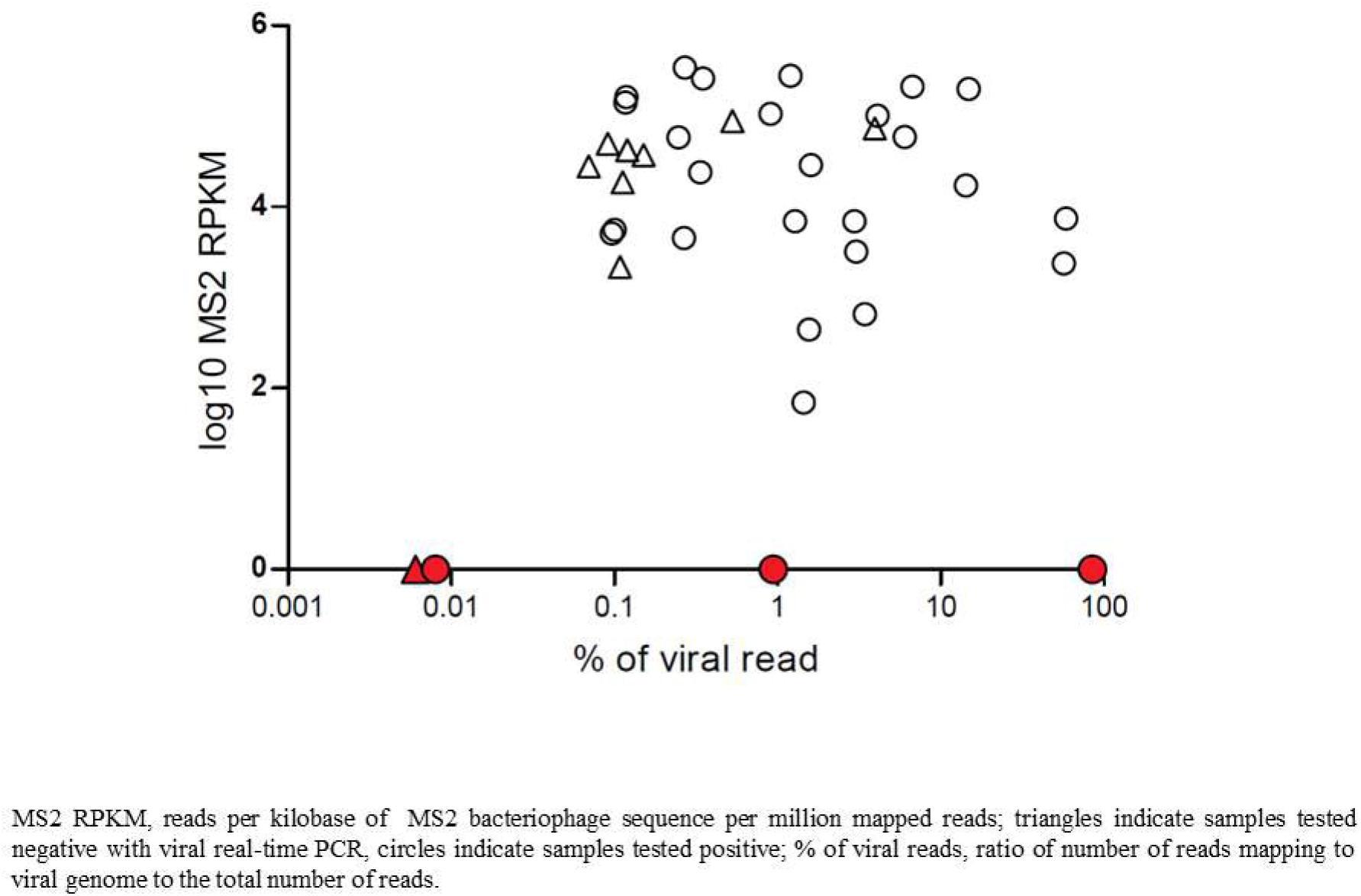
Internal quality control detection after metagenomic analysis of the respiratory samples selected. Distribution of normalized read counts (RPKM) for MS2 bacteriophage (internal quality control, IQC) depending on the proportion of viral reads for the 37 respiratory samples selected. MS2 RPKM was determined after subtracting of NTC MS2 RPKM. IQC was not detected for 4/37 samples (represented in red); among them 3 samples were tested positive with viral real-time PCR.

The 4 samples that did not pass IQCT2 included one sputum that was previously tested negative using real-time PCR (sample # 37), one HCoV positive sputum (sample # 11, Ct = 32), one HBoV positive nasopharyngeal swab (sample # 19, Ct = 30), and one nasopharyngeal aspirate tested positive for HBoV and CMV (sample # 23, Ct = 15 and 31, respectively). For sample # 37 and sample # 19, none of the real-time PCR targeted viruses were detected after bioinformatic analysis. For sample # 19, we sequenced a replicate which similarly failed both IQC and HBoV detection. We could not test any replicate for sample # 37 owing to insufficient quantity. Viral metagenomics results for sample # 23 were validated as viral reads represented 85.2% (9,489,578/11,144,324) of the total reads generated (Fig. 3). For sample # 11, the number of reads mapping to HCoV was 9/5,125,947 with a HCoV genome coverage of 0.2%. Results were therefore not validated for this sample. Overall, mNGS results were validated for 34/37 samples including 26/28 positive samples and 8/9 negative samples.

### Metagenomic workflow evaluation according to viral genome type

The evaluation of the metagenomic workflow was performed using the 26 previously validated respiratory samples tested positive with viral real-time PCR targeting a representative panel of DNA and RNA viruses. For all 26 samples tested, viral metagenomic sequencing allowed the identification of the 36/36 viral genotypes matching targeted PCR results and on-target viral genome coverage ranged from 0.6 to 100% (median = 67.7%). For these 36 targeted viruses, the real-time PCR Ct values ranged from 15 to 37 Ct (median = 28 Ct). The six multiple viral infections involving from 2 to 4 different viruses were also fully characterized (Table 1). For sample # 25 (sample tested positive for 2 DNA viruses and 2 RNA viruses using real-time PCR), mNGS results were cross-checked on a duplicate which reported RPKM deviations lower than 0.5 log for each targeted virus (mNGS results for the 2 replicates are summarized in Additional file 3). Regarding mNGS results obtained from the 8 negative samples validated with IQC, no clinically relevant virus was detected. A strong correlation between mNGS and real-time PCR results was obtained for each viral genome type (R^2^ ranged from 0.72 for linear ssRNA viruses to 0.98 for linear ssDNA viruses, Fig. 4a). Normalized read counts were significantly lower for linear dsDNA viruses than for other viral genome types (Fig. 4b).

**Table 1.** Metagenomic NGS results for the validated respiratory samples tested positive with viral real-time PCR.

| Sample No. | Real-time PCR Ct values |  | Viral genome type | mNGS results for targeted viruses<sup>a</sup> |  |  |  |
| --- | --- | --- | --- | --- | --- | --- | --- |
|  |  |  |  | Identification | No. of reads | RPKM | Coverage(%) |
| 1 | HRV/EV | 25 | linear ssRNA | HRV-A19 | 13,061 | 5.5E+06 | 97.6 |
| 2 |  | 24 |  | HRV-A19 | 29,743 | 8.2E+06 | 98.2 |
| 3 |  | 29 |  | HRV-A63 | 2,672 | 1.4E+06 | 58.1 |
| 4 |  | 34 |  | HRV-A56 | 453 | 1.4E+04 | 75.2 |
| 5 | RSV | 27 |  | RSV-B | 14,218 | 1.9E+06 | 91.2 |
| 6 |  | 36 |  | RSV-A | 187 | 1.5E+03 | 22.0 |
| 7 | MPV | 33 |  | HMPV-A | 44,556 | 9.1E+05 | 100.0 |
| 8 | HCoV | 20 |  | HCoV NL63 | 73,878 | 2.4E+06 | 94.2 |
| 9 |  | 24 |  | HCoV 229E | 19,615 | 1.1E+06 | 99.8 |
| 10 |  | 28 |  | HCoV 229E | 20,666 | 2.4E+05 | 100.0 |
| 12 |  | 36 |  | HCoV NL63 | 1,815 | 1.3E+04 | 9.6 |
| 13 | MV | 23 |  | Measles Virus | 289,019 | 9.1E+06 | 98.1 |
| 14 | IBV | 23 | fragmented ssRNA | Influenza B | 42,212 | 1.1E+06 | 97.2 |
| 15 | IAV | 27 |  | Influenza A(H3N2) | 24,234 | 1.9E+05 | 78.6 |
| 16 |  | 34 |  | Influenza A(H3N2) | 1,559 | 1.9E+04 | 21.2 |
| 17 |  | 35 |  | Influenza A(H3N2) | 258 | 1.8E+03 | 26.5 |
| 18 | HBoV | 24 | linear ssDNA | HBoV-1 | 79,504 | 2.7E+06 | 100.0 |
| 20 | AdV | 17 | linear dsDNA | HAdVC-1 | 245,2476 | 1.6E+07 | 99.8 |
| 21 |  | 36 |  | HAdVD-51 | 18 | 8.0E+01 | 0.6 |
| 22<sup>b</sup> |  | 30 |  | HAdVC-6 | 284 | 1.0E+03 | 6.2 |
|  | HHV-6 | 28 |  | HHV-6B | 18,411 | 1.4E+04 | 54.8 |
| 23<sup>b</sup> | HBoV | 15 | linear ssDNA | HBoV-1 | 9,470,426 | 1.6E+08 | 100.0 |
|  | CMV | 31 | linear dsDNA | CMV | 653 | 2.5E+02 | 5.3 |
| 24<sup>b</sup> | HBoV | 17 | linear ssDNA | HBoV-1 | 7,966,089 | 1.1E+08 | 100 |
|  | MPV | 29 | linear ssRNA | HMPV-A | 10,629 | 5.9E+04 | 95.7 |
| 25<sup>b, c</sup> | AdV | 26 | linear dsDNA | HAdVC-2 | 2,165 | 6.8E+03 | 12.4 |
|  | HPIV | 26 | linear ssRNA | HPIV-3 | 17,576 | 1.3E+05 | 66.7 |
|  | HRV/EV | 34 |  | HRV-C | 446 | 7.0E+03 | 9.2 |
|  | CMV | 27 | linear dsDNA | CMV | 34,577 | 1.7E+04 | 24.8 |
| 26<sup>b</sup> | HRV/EV | 26 | linear ssRNA | HRV-A78 | 114,684 | 1.4E+07 | 99.9 |
|  | AdV | 30 | linear dsDNA | HAdVC-2 | 65 | 1.6E+03 | 9.6 |
|  | RSV | 30 | linear ssRNA | RSV-A | 586 | 3.5E+04 | 68.7 |
| 27<sup>b</sup> | AdV | 32 | linear dsDNA | HAdVC-2 | 24 | 1.3E+02 | 3.2 |
|  | HPIV | 37 | linear ssRNA | HPIV-2 | 50 | 6.3E+02 | 2.3 |
| 28<sup>b</sup> | HRV/EV | 31 |  | HRV-A71 | 1,309 | 3.5E+04 | 61.3 |
|  | EBV | 23 | linear dsDNA | EBV | 2,556 | 3.0E+03 | 39.3 |

HRV: human rhinovirus, EV: enterovirus, RSV: respiratory syncytial virus, HCoV: human coronavirus, HMPV: human metapneumovirus, HPIV: human parainfluenza virus, MV: measles virus, HBoV: human bocavirus, AdV: adenovirus, HHV: human herpes virus, CMV: cytomegalovirus, EBV: Epstein-Baar virus, Ct: Cycle threshold, RPKM: reads per kilobase of virus reference sequence per million mapped reads (normalization of the number of reads mapping to a targeted viral genome).
<sup>a</sup>Targeted viruses: viruses detected with real-time PCR.
<sup>b</sup>Multiple viral infections.
<sup>c</sup>Cross-checked on duplicate sample (deviation <0.5 log).

**Fig. 4.**
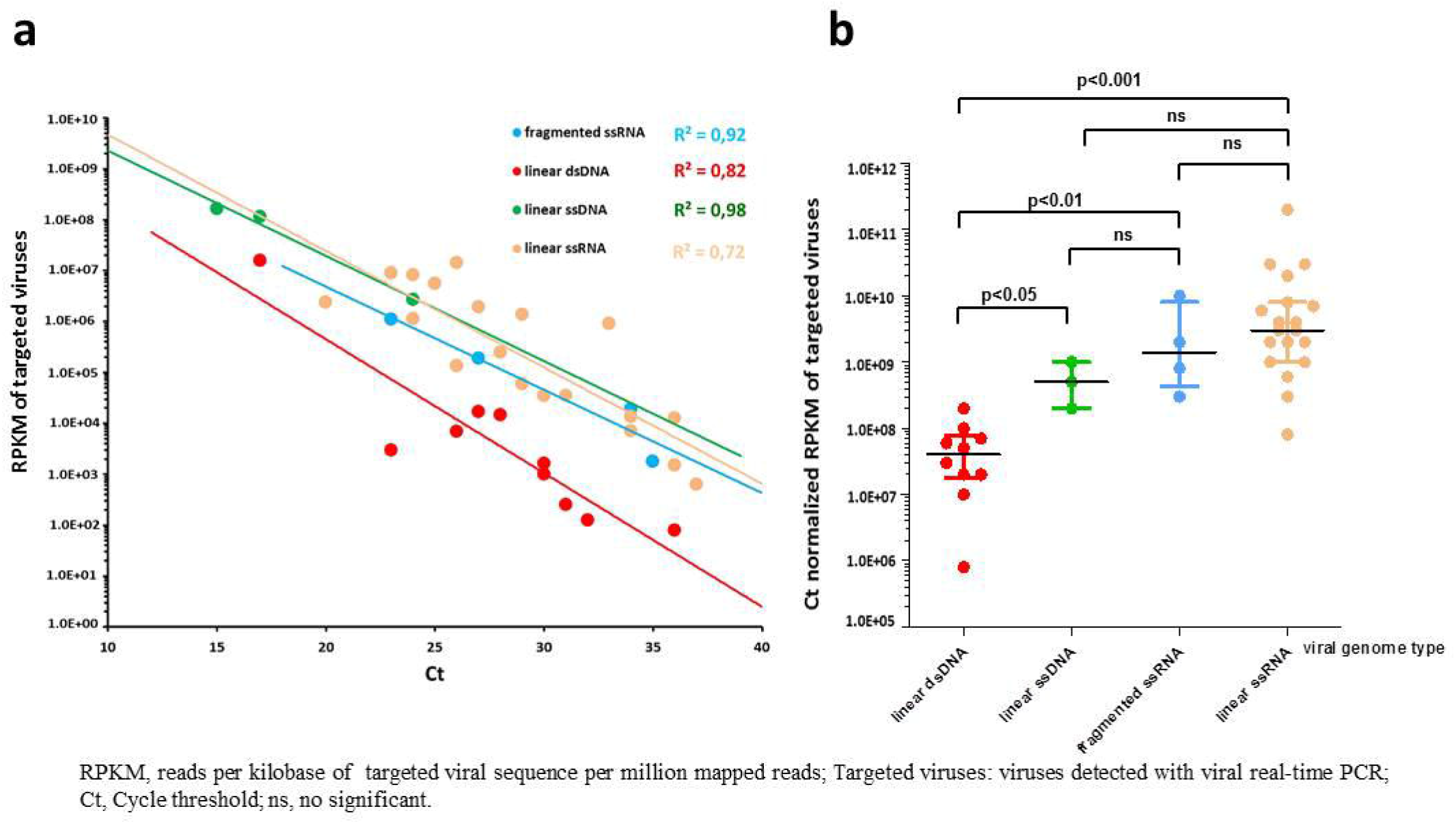
Evaluation of the metagenomic NGS workflow according to the viral genome type. a) Correlation between the results of metagenomic NGS and viral real-time PCR for validated respiratory samples tested positive with viral PCR. Normalized number of reads (RPKM) obtained for targeted virus are displayed against the real-time PCR Ct values for fragmented ssRNA virus (influenza virus) linear dsDNA virus (adenovirus, Epstein-Baar virus, cytomegalovirus, human herpes virus-6) linear ssDNA (human bocavirus) and linear ssRNA (human rhinovirus, respiratory syncytial virus, parainfluenza virus, human coronavirus, human metapneumovirus and measles virus). The correlation coefficients are shown for each viral genome type. b) RPKM normalized by Ct for each viral genome type of validated respiratory samples tested positive with viral PCR. Bars show median and interquartile ranges, p-values calculated with the Mann-Whitney U test are shown.

## Discussion

Over the last few years, a growing number of viral metagenomic protocols have been published but systematic evaluation on clinical respiratory samples and validation by QC is still lacking. In the present study, we describe a process allowing the sensitive detection of both DNA and RNA viruses in a single assay and implemented several QCs to validate the whole metagenomic workflow.

First, IQC was implemented to control the integrity of the reagents, equipment, the presence of inhibitors, and to allow the validation of mNGS results for each sample. The MS2 bacteriophage was selected as IQC for three main reasons; firstly MS2 is widely used as IQC during viral real-time PCR assays to control both extraction and inhibition [24], secondly, an RNA virus was required to control the random reverse transcription and second strand synthesis steps, and thirdly MS2 is a ssRNA virus with a small genome (3,569-bp) that is perfectly characterized and therefore can be easily detected after bioinformatic analysis without the need for extensive NGS reads. The use of MS2 as an IQC has been previously reported for metagenomic analysis of cerebrospinal fluid specimens [25]. In another metagenomic study, RNA of MS2 was included after extraction as an IQC but the use of purified RNA does not validate the viral enrichment step [26]. In the protocol described herein, whole MS2 virions were added to each clinical sample from the beginning of the workflow. QCT1 was implemented to control the first steps of the process and to avoid unnecessary library preparation when these steps fail. At the end of the workflow, QCT2 was able to invalidate 2 samples as neither MS2 nor viruses causing ARIs were significantly detected after metagenomic analysis while routine PCR screenings detected a HBoV and a HCoV. The re-testing of these 2 samples found the same findings suggesting an inhibition or a competition issue during the process. Without the use of IQC, these samples would have been mistakenly classified as false negatives by mNGS. However, the expected competition between viruses and MS2 during the process could lead to a non-detection of IQC reads in case of high viral load. Thus, the interpretation of IQC results should consider the proportion of viral reads of each sample. Although not observed, IQC reads may also be reduced in samples with a greater numbers of patient cells which may affect the sensitivity of the assay [25].

In addition to IQC, we implemented negative external control because contamination issues are frequently reported in metagenomic studies and may lead to misinterpretation in clinical practice [17]. mNGS reads in this negative control were mainly composed of bacterial reads. However, viral reads (mainly derived from prokaryote viruses) were also detected which could be present in reagents (“kitome”) or may represent laboratory contaminants or bleed-over contaminations from highly positive samples within the batch. Such contamination was observed in the present study from the highly positive HBoV sample (sample # 23, Ct=15) which contaminated the NTC (HBoV: 185 reads, RPKM = 1.1E+04 RPKM). In the clinical setting, subtracting NTC viral reads prior to interpretation of each sample result is therefore required.

To evaluate the workflow, clinical respiratory samples tested for a representative panel of DNA and RNA viruses using real-time PCR were selected. This workflow is based on a previous publication where a single protocol had been specifically developed for stool specimens and evaluated on mock communities containing high concentrations of spiked viruses [21]. Interestingly, 6 multiple viral infections involving both DNA and RNA viruses were fully characterized highlighting the power of our mNGS approach as a universal method for virus characterization despite the lack of common viral sequence. In addition to viruses targeted by PCR, viral reads deriving from the commensal virome, including viruses from the *Anelloviridae* family, were generated both in PCR negative and positive samples but not in the NTC.

Regarding the sensitivity of the mNGS approach, a wide range of semi-quantitative real-time PCR Ct values was covered. Thorburn *et al*., compared mNGS to conventional real-time PCR for the detection of RNA viruses on nasopharyngeal swabs and reported a detection cut-off of 32 Ct for the mNGS approach [27]. Our workflow allowed the characterization of both DNA and RNA viruses up to a semi-quantitative real-time PCR Ct value of 36 which is considered to be a low viral load. A major critical point in viral metagenomics is to reduce host and bacterial components. In comparison with similar studies, viral reads herein were highly represented (mean = 7.4%); for example, a study on 16 nasopharyngeal aspirates tested positive with viral PCR assays found a mean of 0.05% of viral reads [12]. In addition, a strong correlation between results of mNGS and conventional real-time PCR was obtained by regrouping viruses according to their genome types. Similar findings were reported elsewhere, suggesting that mNGS results could be used for semi-quantitative measurement of the viral load in clinical samples [3–5,28]. A lower RPKM values for dsDNA viruses compared to the other viral genome types were noticed. As previously described for EBV and CMV, the necessary use of DNase to reduce host contamination may affect these fragile large dsDNA viruses [9,10]. As the detection limit of mNGS analysis is mainly dependent on viral load and total number of reads per sample, this effect could be overcome by increasing sequencing depth; however, we chose to limit the costs of the workflow.

The reagent cost of this mNGS approach is relatively low and was estimated to ∼€150 thanks to our viral enrichment process and the amplification method using a commercial kit which is diluted 5-fold [21]. The use of a universal workflow for both DNA and RNA viruses also reduces the reagent cost compared with metagenomic protocols targeting DNA and RNA viruses separately. In contrast, targeted NGS of specific viruses following their specific amplification by PCR can be up to 2 times cheaper based on our experience (e.g. influenza virus sequencing [29]. Due to several limitations, including its cost and a long turnaround time, viral metagenomics is currently considered to be a second-line approach and is not used as a primary routine diagnostic tool. However, with the improvement of sequencing technologies allowing real-time sequencing such as MinION sequencers (Oxford nanopore, Oxford, United Kingdom), it could be envisioned that mNGS will gradually be used for primary diagnosis in the mid-term. In case of high viral load and sufficient DNA input after amplification our workflow might be used with a MinION sequencer.

The approach described in this preliminary work is crucial to bring standardization for the routine clinical use of mNGS process within a reasonable timeframe. Further evaluation studies with a greater number of samples are urgently needed to establish IQC cut-off according to the number of viral, human and bacterial reads, and to define the performance of the workflow, including repeatability, reproducibility, as well as the detection limit for each virus. In addition, improvement of the bioinformatics pipeline are being explored, including implementation of threshold regarding genome coverage pattern [25], but their impact on performance of the workflow has to be established.

## Conclusion

The potential of mNGS is very promising but several factors such as inhibition, competition, and contamination can lead to a dramatic misinterpretation in the clinical setting. Herein, we provide an efficient and easy to use mNGS workflow including quality controls successfully evaluated for the comprehensive characterization of a broad and representative panel of DNA and RNA viruses in various types of clinical respiratory samples.

## Abbreviations

NGS: Next-Generation Sequencing
mNGS: metagenomic Next-Generation Sequencing
ARIs: Acute Respiratory Infections
PCR: Polymerase Chain Reaction
QC: quality controls
HCL: Hospices Civils de Lyon
IQC: Internal Quality Control
MS2: MS2 bacteriophage
EQC: External Quality Control
NTC: No-Template Control, QCT Quality Control Testing
Ct: Cycle threshold
RPKM: reads per kilobase of virus reference sequence per million mapped reads

## Ethics approval and consent to participate

This single center retrospective study received approval from HCL board of the French data protection authority (*Commission Nationale de l’Informatique et des Libertés*) and is registered with the national data protection agency (number 17–024). Respiratory samples were collected for regular disease management during hospital stay and no additional samples were taken for this study. In accordance with French legislation relating to this type a study, a written informed consent from participants was not required for the use of de-identified collected clinical samples (Bioethics law number 2004–800 of August 6, 2004). During their hospitalization in the HCL, patients are made aware that their de-identified data including clinical samples may be used for research purposes, and they can opt out if they object to the use of their data.

## Consent for publication

Not applicable.

## Availability of data and materials

Data generated during this study are included in supplementary information files. Sequencing datasets used and/or analysed during the current study are available from the corresponding author on reasonable request.

## Authors’ contributions

AB, LJ, FM, KB SA conceived the study. AB, MP, LB, VC performed the sample preparations and sequencing. LJ, GO, GV performed bioinformatic analysis. LJ is the guarantor for the NGS data. YG, MV, IS, BL, SV, FM are the guarantor for clinical data and sample collection. AB was the main writer of the manuscript. All authors reviewed and approved the final version of the manuscript.

## Acknowledgments

We thank Audrey Guichard, Gwendolyne Burfin, Delphine Falcon and Cecile Darley for their technical assistance as well as Philip Robinson (DRCI, Hospices Civils de Lyon) for his excellent help in manuscript preparation. Part of these data has been presented at the International Conference of Clinical Metagenomic held in Geneva in October 2017.

## Titles and legends for additional files

**Additional file 1. (Table.xls) Summary of clinical samples and metagenomic NGS information**.

**Additional file 2. (Table.xls) Quality control testing results**.

QCT1 corresponds to MS2 bacteriophage molecular detection with commercial real-time PCR assay. QCT2 corresponds to control by sequencing metrics (number of MS2 reads normalized with RPKM ratio and MS2 genome coverage). MS2 RPKM for the 37 selected clinical samples was determined after subtracting of NTC MS2 RPKM.

**Additional file 3. (Figure.ppt) Metagenomic NGS results for duplicates of sample # 25**.

Sample # 25 corresponds to a clinical respiratory sample tested positive for 2 DNA viruses (adenovirus, cytomegalovirus) and 2 RNA viruses (human parainfluenza virus, human rhinovirus) using real-time PCR. This sample was analyzed twice using our single metagenomic workflow (replicate 1 and replicate 2). a) Pie charts show classification of reads into human, bacteria, viruses, fungi, archea and unknown categories (unassigned reads). b) Normalized read counts (RPKM) for each targeted virus (viruses detected with real-time PCR) and for internal quality control (MS2 bacteriophage). c) Coverage plot of targeted viral genomes and internal quality control (MS2 bacteriophage). Sequencing reads were mapped on ezVIR viral database that identified human adenovirus C-2 (accession number: KF268130.1), cytomegalovirus (accession number: GQ396662.1), human parainfluenza virus 3 (accession number: KF687321.1), human rhinovirus C (accession number: JF317014.1) and MS2 bacteriophage (accession number: NC_001417.2).

